# Age-specific and Sex-specific Nectar and Pollen Use by a Butterfly Pollinator

**DOI:** 10.1101/2022.05.19.492749

**Authors:** Carol L. Boggs, Vikram Iyengar

## Abstract

Age-specific patterns of nectar and pollen use by insect pollinators may reflect behavioral or physiological changes over the insect’s lifespan, and may also influence flower visitation rate. Studying *Heliconius charithonia* (Lepidoptera: Nymphalidae) and *Psiguria umbrosa* (Cucurbitaceae), we showed that honey-water (as a nectar substitute) intake increases over the first ten to twelve days of female’s life, while decreasing over the same age period for males, when individuals were fed *ad libitum*. Intake then remains stable at least through 28 days of age. Mean intake is higher for females than for males, and is not significantly affected by body size. Intake patterns for honey-water and pollen did not change with age in a similar manner. Pollen feeding increased significantly with age for both sexes, with females showing a two week delay in the increase when studied in a free-flying greenhouse population with competition for pollen. Under both *ad libitum* and pollen competition conditions, females collected more pollen than did males. Body size did not significantly affect the amount of pollen collected for either sex. Within each sex, butterflies with *ad libitum* pollen collected more pollen than those in a free-flying greenhouse population with more restricted pollen availability. These results suggest that two resources obtained from the same source, pollen and nectar, are not treated identically by the insect pollinator, and that the demography of the insect population may affect flower visitation rates in some cases. Further, foraging patterns for pollen and nectar are likely driven by physiological demand for different resource types.

## Introduction

Flower visitation rates by insect pollinators are a result of both plant and pollinator characteristics (e.g., Boggs 1987). Important plant traits include flower morphology (e.g., Grant 1949, Stebbins 1970, Barrows 1976, Stuessy et al. 1986), physical cues such as scent and color (e.g., Kevan 1978), temporal patterns of reward availability (e.g., Pleasants 1983, Devlin and Stephenson 1985, Zimmerman 1988, Rotenberry 1990), distances among flowers (e.g. Levin and Kerster 1969), and spatial distribution of rewards among flowers (e.g., Pleasants and Zimmerman 1979, Ott et al. 1985, Waser and Mitchell 1990). Total reward availability within a given area can also affect visitation rates to individual flowers (e.g., Augspurger 1980, Murawski 1987, Thomson 1987, Zimmerman 1988). Insect traits affecting flower visitation include mouthpart morphology (e.g., Emmel 1971, Gilbert and Singer 1975, Barrows 1976, Gilbert 1981), ability to thermoregulate (e.g., Kingsolver 1983), and hence forage, under extreme ambient temperature conditions, flight distance between flower visits (independent of flower density) (e.g., Schmitt 1980, Waser 1982, Zimmerman 1988, Rathcke 1992), ability to learn flower location or flower type with consistent rewards (e.g., Levin and Berube 1972, Schemske 1976, Menzel 1990), ability to learn the location of parasitized flowers (Murawski 1987) and energetic costs associated with foraging (e.g., Heinrich 1975, Kammer and Heinrich 1978). The importance of these traits to insect energy balance and both pollen placement efficiency and vector pollinating efficiency (*sensu* Inouye et al. 1994) is well-documented in the literature.

Changes in pollinator behavior with age could also influence flower visitation rates. Changes in learning ability or propensity with age could affect flower choice (e.g., Lewis 1989). Physiological requirements may change with age and vary between the sexes. Foraging costs may change with age, for example through changes in flight ability as wings fray with use. Cartar (1992) showed that wing damage in bees may result in increased mortality or a halt to foraging by experimental individuals. Changes in nectar intake rates with age (Boggs 1988) may also affect foraging costs. The risk of mortality that an individual is willing to incur for a given nutritional reward may change with reproductive value, and hence age, leading to changes in optimal foraging patterns (Wilson 1985, Engen and Stenseth 1989).

In spite of evidence that insect age may be an important factor affecting flower visitation rate, there are only a few direct measures of the effect of age on nectar or pollen intake, or on pollen transfer ability. Nectar intake varies with age and sex in at least one butterfly species, *Speyeria mormonia*, under laboratory conditions (Boggs and Ross 1993). Females imbibe more nectar than do males, and nectar intake decreases with age in both sexes. Pollen collection and feeding also vary with age and sex in another butterfly genus, *Heliconius*. Females collect more pollen than do males, and pollen collecting increases with age (Boggs et al. 1981, Boggs 1990). Within social hymenoptera, pollen and nectar collection also varies among castes (Wilson 1985).

Some insect pollinators use both nectar and pollen from the same plant species. However, age- and sex-specific patterns of resource use need not be similar for both pollen and nectar. Further, efficiency of pollen transfer may vary, depending on whether the insect is foraging for nectar or pollen, particularly if pollen feeding involves retention of pollen on the proboscis or other body parts that may contact a stigma. The degree of similarity of age- and sex-specific patterns of nectar and pollen use could thus affect how tightly nectar use is coupled to pollen transfer, potentially affecting the strength of selection on nectar production rates. Likewise, if pollen and nectar use do not change in synchrony among insect ages and sexes, the effectiveness of an individual insect as a pollinator may change over time. Further, the effectiveness of the insect population as pollinators may vary through time, depending on changes in the population age and sex structure.

Here we examine the age- and sex-specific patterns of nectar and pollen use by one species of butterfly, *Heliconius charithonia* (Nymphalidae). Members of this butterfly genus are pollinators of *Psiguria* (Cucurbitaceae) and other species in some neotropical localities (Murawski 1987).

## Methods

### Study system

*Heliconius charithonia* is a neotropical butterfly which feeds as an adult on both nectar and pollen, including that from *Psiguria umbrosa*. Pollen is collected shortly after anther dehiscence, and held as a mass on the proboscis. A fluid is exuded, and mixed over a period of several hours with the pollen mass, eventually causing the pollen to begin to germinate. Germinating pollen releases enzymes, amino acids, peptides, nucleotides, etc., into the fluid. The enriched fluid is then imbibed by the butterfly, and the pollen mass is eventually sloughed off the proboscis after several hours (Gilbert 1973). Pollen feeding is necessary for normal lifespan and egg production by *Heliconius* species in general (Gilbert 1973, Dunlap-Pianka et al. 1977).

*P. umbrosa* is a neotropical vine with male and female flowers. Individual plants can produce either male or female flowers, or both sequentially or simultaneously (Condon & Gilbert 1988; Boggs pers. obs.). Male flowers far outnumber female flowers (Boggs pers. obs.)

### Experimental Designs

#### Honey-water Use

Ten male and ten female *H. charithonia* were maintained in 1m x 1m x 1.5m screen cages inside a greenhouse from adult emergence to 28 days of age. The cages contained more than one *P. umbrosa* flower per butterfly; one flower produces as much nectar per day as an individual butterfly will eat (Iyengar, unpublished). Females were mated on the day of emergence and had access to larval host plants for oviposition, but males were not given opportunities to mate. At adult emergence, we noted length of the forewing (a measure of body size), and individually numbered butterflies on their wings using a felt marker.

Intake of honey-water, as a nectar substitute, was measured every third day until 28 days of age, beginning on the third day of adult life. Individual butterflies to be tested were isolated from nectar plants overnight prior to test, and throughout the day of the test. We fed the butterfly a 1:3 (v:v) honey:water solution, using a 25 µl Hamilton syringe to measure the amount imbibed. Butterflies were fed until they rejected the solution three times. The total amount imbibed was recorded. Feeding was done twice on each experimental day, at 9:00 and 16:00.

Refractometer measurements of both *P. umbrosa* nectar and our honey-water solution gave readings of 18-20% (Boggs, unpublished). Although the exact composition of nectar may differ from the honey-water solution, the percent dissolved solids was thus similar, which should lead to similar feeding rates and costs (Kingsolver & Daniel 1979, May 1985, Pivnick & McNeil 1985, Boggs 1988, Daniel et al. 1988)

#### Pollen Use

We examined pollen use by *H. charithonia* under two conditions. The first allowed *ad libitum* access to pollen. The second was done on a free-flying greenhouse population, with limited access to pollen and hence competition among individuals for pollen.

For the first study, with *ad libitum* pollen access, 19 males were maintained in 1m x 1m x 1.5m screen cages inside a greenhouse from adult emergence until death or 45 days of age, whichever came first. 17 females were maintained individually in 1m x 0.5m x 1.3m screen cages. All cages contained one or more *P. umbrosa* flower per butterfly; one flower produces more pollen than one butterfly will eat per day (Boggs, unpub. observations). Females were mated on the day of adult emergence, with three males available as mates, and had access to larval host plants for oviposition. Males were not given the opportunity to mate. Forewing length was recorded for all males and 7 females, as an indicator of body size.

For the second study, with restricted pollen access, *H. charithonia* were maintained as a free-flying greenhouse colony. The greenhouse contained larval host plants and *P. umbrosa* as an adult nectar and pollen source, as well as a population of *H. cydno*, which compete with *H. charithonia* for pollen and nectar. The number of *P. umbrosa* flowers per butterfly was not recorded, but was less than one per butterfly. Males could mate freely, and females could oviposit freely. (Females only mate once in this species, while males may mate several times (Boggs 1979).) Under these conditions, individuals’ lifespans, mating habits, and pollen collecting habits were similar to those seen in the field (Boggs 1979). Data were used from 46 males and 36 females, over the first 45 days of adult life or until death, whichever occurred first.

For both studies, individual butterflies were numbered on the day of adult emergence with a felt-tip marker. For each individual, pollen load sizes were visually scored daily on a linear scale of 0-3 used by Boggs et al. (1981), where 0 = no pollen and 3 = ∼3000 pollen grains. Pollen load scoring occurred between 9:00 and 11:00, as pollen is collected in the morning.

## Results

### Honey-water Use

Daily *ad libitum* intake of honey-water differed among individual butterflies (table 1). Averaging among individuals’ mean intake, females imbibed 53±5 (x^-^±s.d.) µl/day, significantly greater than the 42±5 µl/day eaten by males (t = 4.85, 18 d.f., P<0.001). Winglength, as an indicator of body size, had no effect on mean intake for either sex (males: F_1,8_ = 1.11, ns; females: F_1,8_ = 2.21, ns).

**Table 1.**
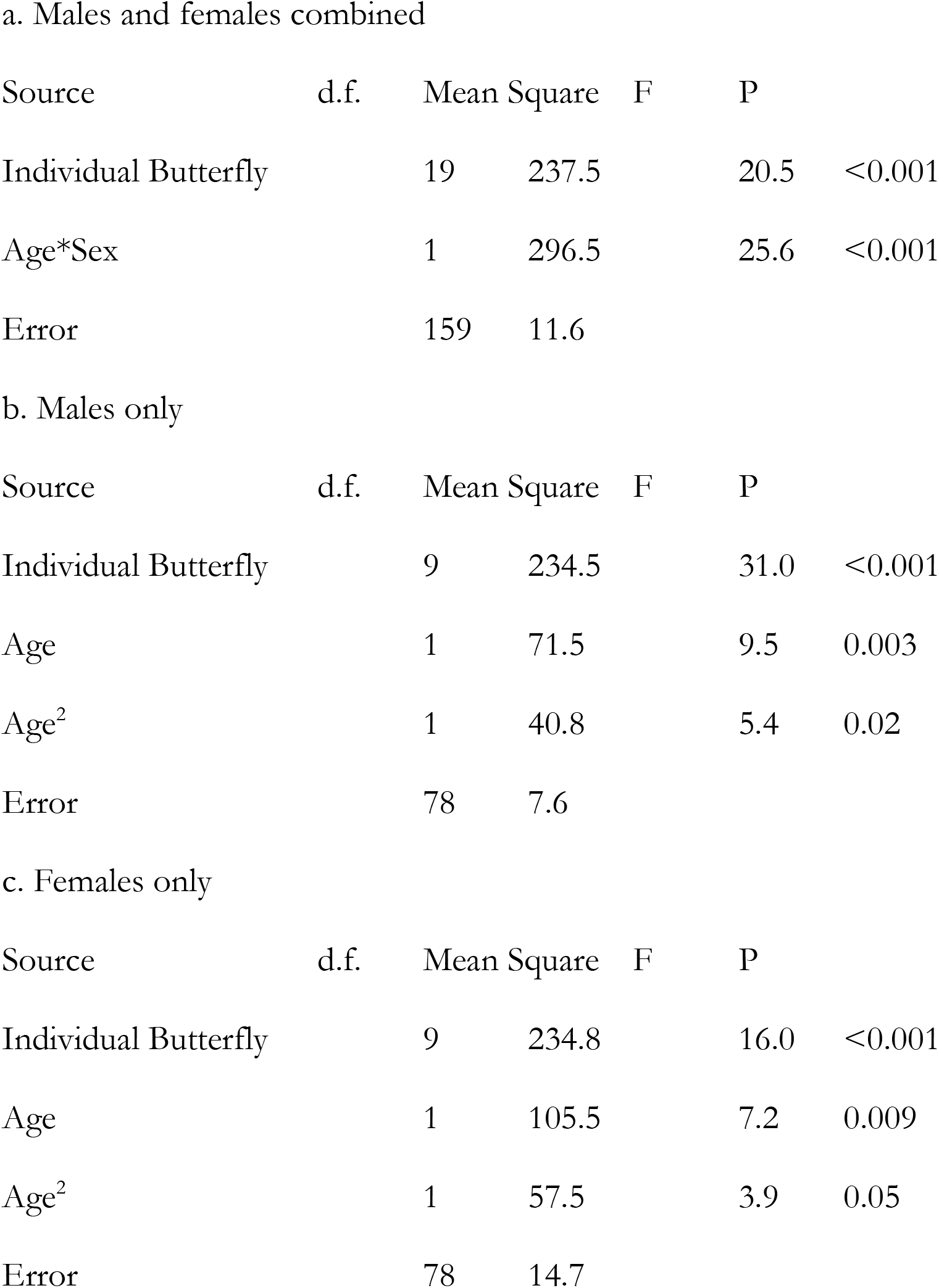
Effects of age and sex on amount of honey-water (a nectar substitute) imbibed daily for the first 28 days of adult life by *H. charithonia* under *ad libitum* conditions.

The age-specific pattern of honey-water intake differed between the two sexes (table 1; fig. 1). Regressions of honey-water intake against age, with individual butterfly as a category variable, showed that intake decreases with age, and then flattens out for males, while increasing with age and then flattening out for females (table 1; fig. 1). In both cases, the quadratic term was significant. Intake stabilized around 12 days of age for both sexes.

**Fig 1.**
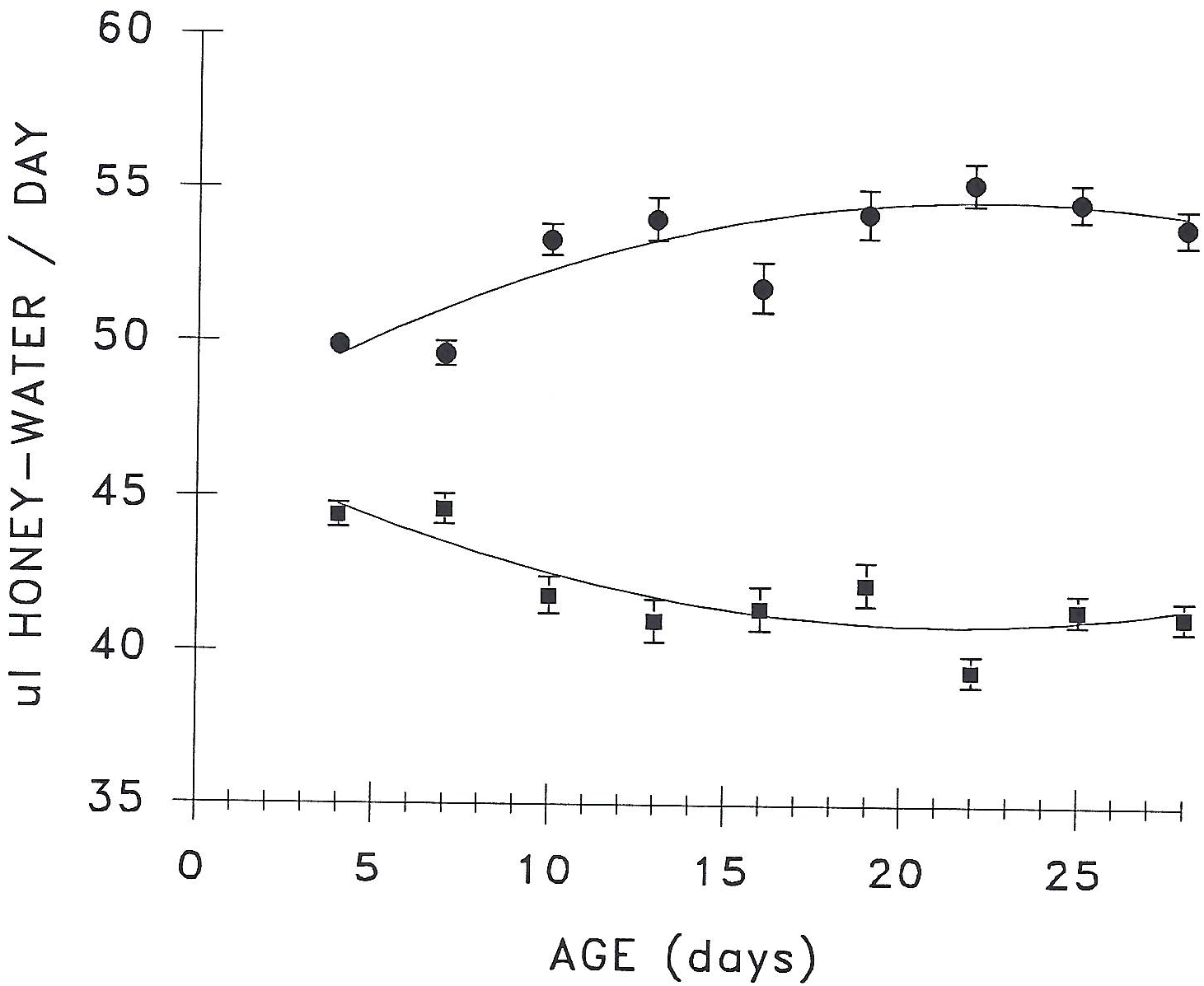
Mean and standard error of age-specific intake of honey-water (as a nectar substitute) by *H. charithonia* fed *ad libitum*. ◼= males, • = females. Lines are regressions of intake against age, with individual butterfly included in the analysis (see table 1 for significance tests). Males: y = 46.77 - 0.55x + 0.01x^2^. Females: y = 47.09 + 0.67x - 0.02x^2^.

### Pollen Use

Daily pollen use differed among individual butterflies (table 2). Averaging individuals’ lifetime mean pollen load scores and with lifespan as a covariate, females had significantly greater intake than males for both the *ad libitum* and free-flying population treatments, and *ad libitum* intake was higher than intake in the free-flying population treatment for both sexes (table 3). Winglength had no effect on mean intake for *ad libitum* fed individuals of either sex, with lifespan as a covariate (males: t = -0.37, 17 df, ns; females: t = 0.742, 5 df, ns). Winglength was not measured for the free-flying population data set.

**Table 2.** Effects of age on daily pollen load score during the first 45 days of adult life for *H. charithonia*. a. Males, maintained as free-flying greenhouse populationc. Females, maintained as free-flying greenhouse population Source d.f. Mean Square F P

| Source | d.f. | Mean Square | F | P |
| --- | --- | --- | --- | --- |
| Individual Butterfly | 45 | 0.1 | 2.9 | <0.001 |
| Age | 1 | 0.5 | 13.5 | <0.001 |
| Age<sup>2</sup> | 1 | 0.3 | 9.0 | 0.003 |
| Error | 1172 | 0.04 |  |  |

b. Males, fed *ad libitum*
| Source | d.f. | Mean Square | F | P |
| --- | --- | --- | --- | --- |
| Individual Butterfly | 18 | 0.7 | 4.5 | <0.001 |
| Age | 1 | 4.6 | 30.8 | <0.001 |
| Age<sup>2</sup> | 1 | 3.1 | 20.7 | <0.001 |
| Age<sup>3</sup> | 1 | 2.4 | 16.1 | <0.001 |
| Error | 570 | 0.1 |  |  |

c. Females, maintained as free-flying greenhouse population
| Source | d.f. | Mean Square | F | P |
| --- | --- | --- | --- | --- |
| Individual Butterfly | 35 | 0.5 | 4.9 | <0.001 |
| Age<sup>2</sup> | 1 | 6.0 | 62.3 | <0.001 |
| Age<sup>3</sup> | 1 | 3.9 | 40.3 | <0.001 |
| Error | 895 | 0.1 |  |  |

d. Females, fed *ad libitum*
| Source | d.f. | Mean Square | F | P |
| --- | --- | --- | --- | --- |
| Individual Butterfly | 16 | 6.3 | 11.8 | <0.001 |
| Age | 1 | 10.2 | 19.2 | <0.001 |
| Age<sup>2</sup> | 1 | 5.6 | 10.5 | 0.001 |
| Error | 447 | 0.5 |  |  |

**Table 3.**
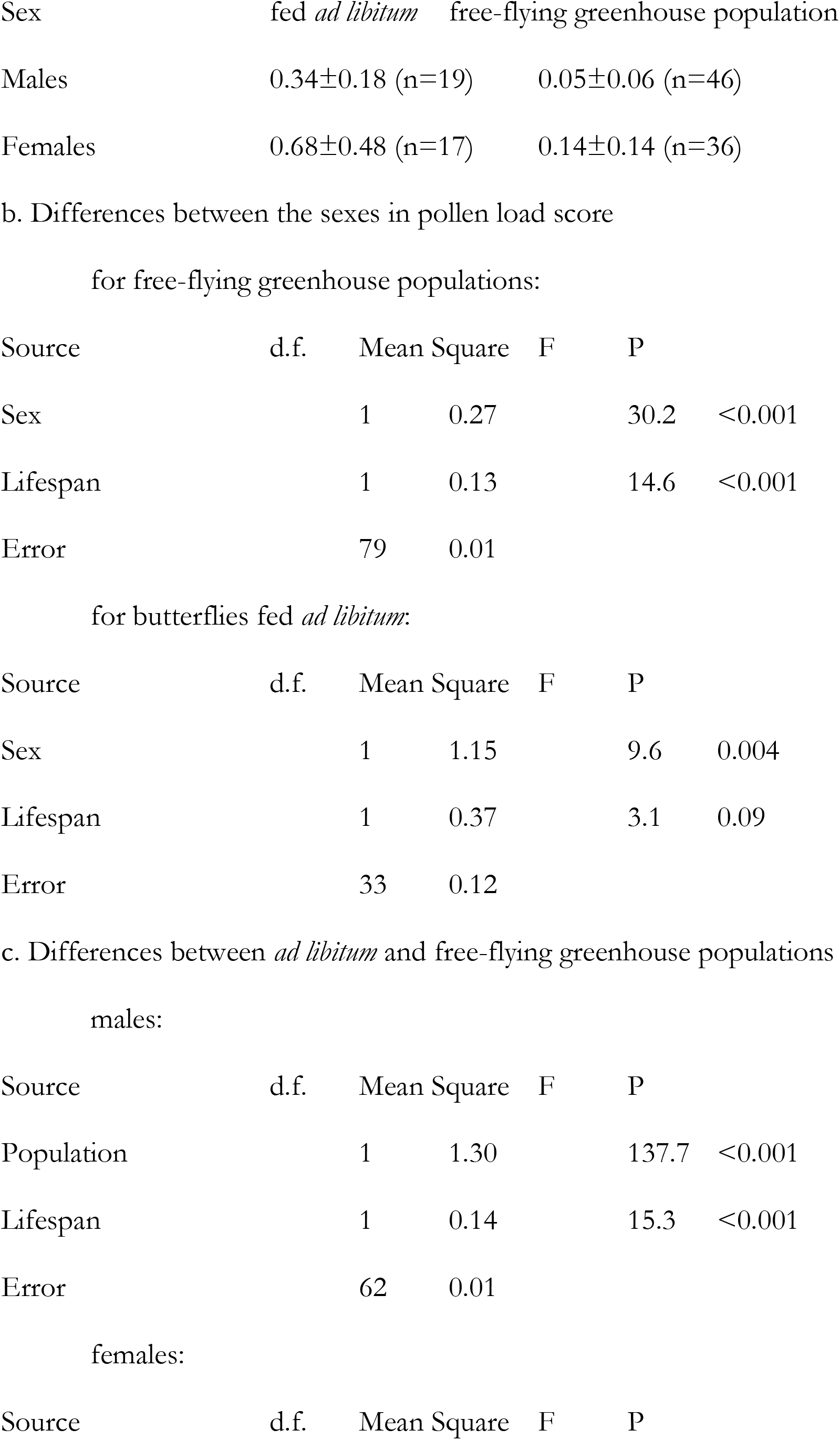
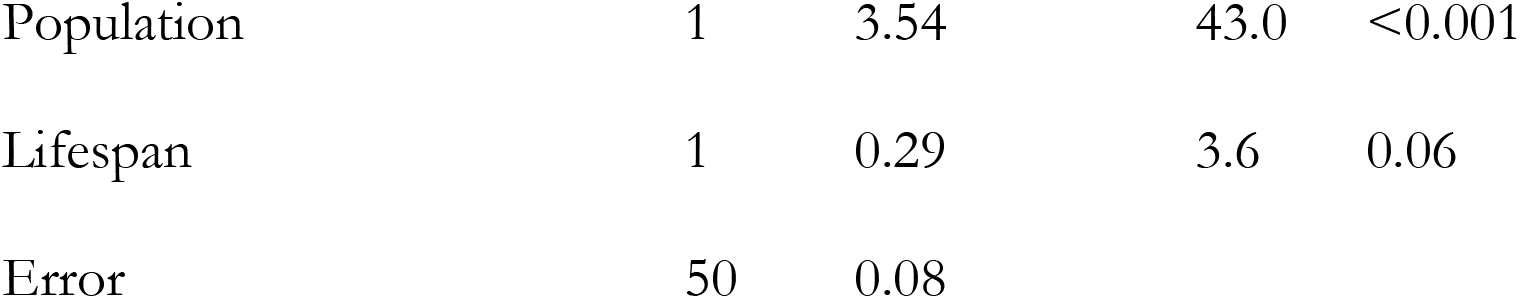
Daily pollen load score for *H. charithonia* over the first 45 days of adult life.

| Sex | fed *ad libitum* | free-flying greenhouse population |
| --- | --- | --- |
| Males | 0.34±0.18 (n=19) | 0.05±0.06 (n=46) |
| Females | 0.68±0.48 (n=17) | 0.14±0.14 (n=36) |

b. Differences between the sexes in pollen load score for free-flying greenhouse populations:
| Source | d.f. | Mean Square | F | P |
| --- | --- | --- | --- | --- |
| Sex | 1 | 0.27 | 30.2 | <0.001 |
| Lifespan | 1 | 0.13 | 14.6 | <0.001 |
| Error | 79 | 0.01 |  |  |

for butterflies fed *ad libitum*:
| Source | d.f. | Mean Square | F | P |
| --- | --- | --- | --- | --- |
| Sex | 1 | 1.15 | 9.6 | 0.004 |
| Lifespan | 1 | 0.37 | 3.1 | 0.09 |
| Error | 33 | 0.12 |  |  |

c. Differences between *ad libitum* and free-flying greenhouse populations males:
| Source | d.f. | Mean Square | F | P |
| --- | --- | --- | --- | --- |
| Population | 1 | 1.30 | 137.7 | <0.001 |
| Lifespan | 1 | 0.14 | 15.3 | <0.001 |
| Error | 62 | 0.01 |  |  |

females:
| Source | d.f. | Mean Square | F | P |
| --- | --- | --- | --- | --- |
| Population | 1 | 3.54 | 43.0 | <0.001 |
| Lifespan | 1 | 0.29 | 3.6 | 0.06 |
| Error | 50 | 0.08 |  |  |

Regressions of pollen load score against age, with individual butterfly as a category variable, showed that male pollen load scores increased significantly with age, and then roughly levelled off after about 11-15 days for both *ad libitum* and greenhouse populations (fig. 2, table 2). Individuals fed *ad libitum* showed a significant age^3^ effect, with an increase in pollen feeding again around 40 days.

**Fig 2.**
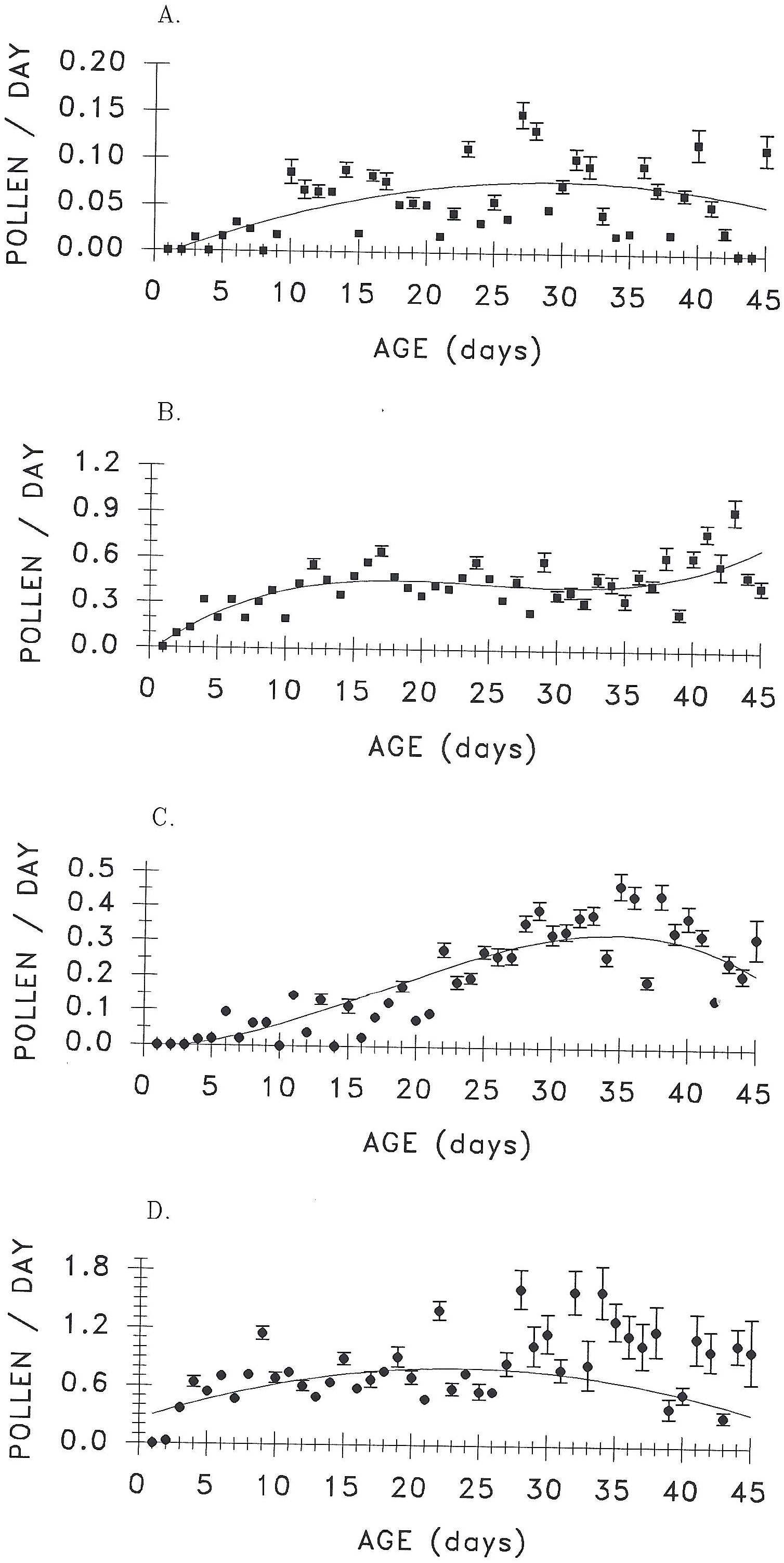
Mean and standard error of age-specific pollen load score for *H. charithonia*. Lines are the regression of pollen load against age, with individual butterfly included in the analysis (see table 2 for significance tests). a. Males in a free-flying greenhouse population. y = -0.009 + 0.006x - 0.0001x^2^ b. Males fed *ad libitum*. y = -0.036 + 0.067x - 0.003x^2^ + 0.00004x^3^. c. Females in a free-flying greenhouse population. y = -0.006 + 0.001x^2^ - 0.00002x^3^. d. Females fed *ad libitum*. y = 0.254 + 0.047x - 0.001x^2^

The age-specific pattern of pollen intake differed between greenhouse and *ad libitum* treatments. For females in the greenhouse population, pollen load scores remained relatively low until about 15-18 days of age, when they began to increase, flattening out about 12 days later at 30 days (fig. 2, table 2). Regression of pollen load score against age, with individual butterfly as a category variable, showed that this pattern was significant. For females fed *ad libitum*, pollen intake increased rapidly with age in a pattern more reminiscent of that for males, levelling off within the first week of life (fig. 2, table 2). Regression of pollen load score against age, with individual butterfly as a category variable, showed that this pattern was significant.

## Discussion

Use of pollen and honey-water (as a nectar substitute) both changed with age. The change in intake was particularly pronounced during the first two weeks of life for nectar feeding by both sexes and for pollen feeding for all but females experiencing competition. For males, nectar use was highest early in life, while pollen use was lowest during that time. Females exhibited an earlier increase in nectar use than in pollen use under pollen competition, but not under *ad libitum* pollen feeding conditions.

We do not know how the pattern of nectar use would differ under competition from that observed when butterflies are fed *ad libitum*. Such an experiment is obviously more difficult to design, as individual nectar intake is more difficult to monitor passively. Nonetheless, differences in pollen collection under *ad libitum* conditions and competition suggest that nectar intake levels could be lower under competition, and could exhibit delays in the female increase in intake and/or more rapid decreases in male intake.

### Plant and butterfly perspectives

Based on the results reported here, from the plant’s perspective, the butterfly population’s age and sex distributions may affect pollen placement and vector pollinating efficiency, assuming that flower visits to collect pollen result in differing pollen transfer efficiency than visits to collect only nectar. Although *Heliconius* have overlapping generations, and population numbers are relatively stable in some areas (Ehrlich and Gilbert 1973), *H. charithonia* is a fairly “weedy” inhabitant of successional areas, and is more likely than many *Heliconius* species to show variation in age structure and density through time within one population.

From the butterfly’s perspective, nectar and pollen are clearly treated as different resources, even though both come from the same source, male flowers. Differences between pollen and nectar intake patterns are likely tied to physiological requirements for carbohydrates and nitrogenous compounds by adults. For example, the delay in pollen feeding seen in the greenhouse female population correlates with the time during which nitrogen-rich spermatophore nutrients donated by males are used by the female in egg production (Boggs 1990), yet sugars are presumably needed to support female flight activity from the outset of adult life. The difference between age-specific pollen feeding habits of females allowed *ad libitum* access to pollen as opposed to those in the free-flying population suggests that when pollen is readily available, females will collect it early in life, perhaps either storing the nutrients gained or delaying use of male-donated nutrients in the spermatophore.

Initial changes in the amount of pollen or nectar eaten took about two weeks from the beginning of life for all male treatments. The consistency in this time block suggests either that there is a fixed behavioral maturation time needed to develop a given feeding behavior fully, or that a nutritional homeostatic state is reached after that time point. Likewise, amount of honey-water imbibed levelled off after about two weeks for females, and the increase in pollen feeding took about two weeks in the greenhouse population, once it began at about age 18-20 days. However, pollen feeding plateaued much more rapidly for females fed pollen *ad libitum*, suggesting that for females and pollen at least, there is no fixed behavioral maturation time of two weeks, but nutritional homeostasis is the most likely cause of levelling off of pollen load scores.

Individual butterflies differed significantly in all treatments for both honey-water and pollen intake. This variation was not due solely to differences in lifespan, or to differences in body size. Fitness effects of such variation, if any, remain to be explored.

### Contrasts with other data sets

The pollen feeding data reported here are consistent with patterns seen previously in field populations of *Heliconius*. For ten *Heliconius* species studied in Costa Rica and Trinidad, Boggs et al. (1981) found that males generally collected less pollen than females, and that pollen loads often increased with age where age was based on wing condition. Further, the slight downward trend in pollen load score seen in the present free-flying greenhouse population near 40 days of age is consistent with a downward trend seen in older age classes for *H. sara* at La Selva, Costa Rica, although not for *H. ethilla* in Trinidad or *H. hecale* at Paloverde, Costa Rica. An *H. charithonia* sample from Santa Rosa, Costa Rica was included in the earlier study; males collected significantly less pollen than females. The change in load size with age was not significant, although the means show the same trend as that seen in the present data.

The difference between the sexes in mean honey-water intake seen here in *H. charithonia* parallels that seen previously for *S. mormonia* (Boggs & Ross, 1993); females imbibed more than males. The lack of effect of body size, as measured by fore wing length, was also consistent between the two species. However, the age-specific patterns differed dramatically between the species. Intake declined with age in both sexes for *S. mormonia*, while either increasing or decreasing to a fairly stable plateau over the same age span in *H. charithonia*. These differences may be tied to differences in the physiology of the two species. Female *H. charithonia* show continual oogenesis, as long as pollen is available (Dunlap-Pianka et al. 1977), whereas *S. mormonia* eclose as adults with a fixed number of oocytes in the ovaries (Boggs 1986). Eggs are thus continually yolked and laid in *H. charithonia*, but the number laid per day in *S. mormonia* declines after about two weeks. Adult glucose sources are used in egg production, at least for *S. mormonia* (Boggs mss). Thus, differences in egg production patterns may result in differences in age-specific requirement for nectar, with a more stable plateau in honey-water intake resulting from a more stable egg production pattern in *H. charithonia*. Nonetheless, this explanation cannot account for the initial differences between the species in honey-water intake, which increased in *H. charithonia* and decreased in *S. mormonia*, since both species’ adults eclose with no mature eggs in the ovaries.

## Conclusions

Foraging patterns for nectar and pollen change with sex and age in *H. charithonia*, as well as with flower availability. The exact mechanisms underlying these changes remain to be determined. However, they are likely driven by physiological demand for different nutrient types. Willingness to accept the risks associated with foraging, or changes associated with behavioral maturation may also play a role.

Age- and sex-specific changes in foraging behavior are likely for other insect pollinators as well. The impact of such changes on the evolution of pollinator/plant associations, as well as on plant population genetic structure, needs to be examined more fully.

## Acknowledgements

We thank A. Brody, D. Inouye and N. Waser for comments on the manuscript.

